# Higher transcriptome stability during aging in long-lived giant mole-rats compared to short-lived rats

**DOI:** 10.1101/251777

**Authors:** Arne Sahm, Martin Bens, Yoshiyuki Henning, Christiane Vole, Marco Groth, Matthias Schwab, Matthias Platzer, Karol Szafranski, Philip Dammann

## Abstract

Many aging-associated physiological changes are known to come up in short‐ and long-lived species with a different trajectory and emerging evidence suggests that large parts of life history trait differences between species are based on inter-species variation in gene expression. Little information is yet available, however, about transcriptome changes during aging when comparing mammals with different lifespans. For this reason, we studied the transcriptomes of five tissues and two age cohorts in two similar sized rodent species with very different lifespans: rat (*Rattus norvegicus*) and giant mole-rat (*Fukomys mechowii*) with maximum lifespans of 3.8 and >20 years, respectively. Our results show that giant mole-rats exhibit higher transcriptome stability during aging than the rat. While well-known aging signatures (e.g. up-regulation of pro-inflammatory genes) were detected in all rat tissues, they showed up only in one giant mole-rat tissue. Furthermore, many differentially expressed genes that were found in both species, were regulated in opposite directions during aging. This suggests that expression changes that cause aging in short-lived species are counteracted in long-lived species. Taken together, transcriptome stability may be one key causal factor of the long life‐ and healthspan of giant mole-rats and maybe of African mole-rats in general.

## Introduction, results, and discussion

Long-lived mammal species have repeatedly been shown to exhibit less age-associated changes in numerous physiological parameters that are assumed to be related to the functional decline during aging than short-lived ones[1-4]. Evidence from recent RNA-seq studies suggest that major parts of the remarkable lifespan diversity amongst mammals are based on inter-species differences in gene expression [5, 6]. However, these studies have focused on the identification of particular genes and pathways that are differently expressed between species with divergent longevities. Whether short-and long-lived species differ regarding the general stability of their transcriptomes has, to the best of our knowledge, never been explored so far.

To address this question, we examined transcriptome changes with age in two similar sized rodent species with different longevities, the laboratory rat (*Rattus norvegicus*) which has a maximum lifespan of 3.8 years [7] and the giant mole-rat (*Fukomys mechowii*), which has a maximum lifespan of >20 years [8]. In giant mole-rats longevity depends strongly on reproductive status, with breeding individuals outliving non-breeders by far [8]. In this study, only non-breeding male individuals were examined. Male non-breeding giant mole-rats have a maximum lifespan of ∼10 years and an average lifespan of ∼6 years, thus still clearly exceeding the life expectancy of the rat [8]. For both species, we performed RNA-seq across five tissue samples (blood, heart, kidney, liver and skin; hereinafter called for simplicity “tissues”) in groups of young and elderly adults, determined differentially expressed genes (DEGs) and searched for enriched functional categories. The analyzed rats had an age of 0.5 (n=5) and 2 (n=4) years, while the giant mole-rats were sampled at mean ages of 1.53 (range 1.3-2.0, n=4-7) and 6.64 (5.5-7.7, n=4-8) years (Table S1). The later time points correspond to an age-associated survival that is about or even below 40% in both species [8, 9]. The earlier time points represent young, yet sexually mature adults and were chosen to be approximately one quarter of the respective later time point.

Despite the fact that both species were compared across a similar range of adult biological age (as derived from the survival probabilities of the cohorts), strikingly, the transcriptomes of the giant mole-rats changed much less than those of the rats. In four of five tissues the number of DEGs in the giant mole-rat represented only a small fraction of the respective numbers in the rat (0.6-19%, Fig. 1, tables S2-11). Only in blood, the number of DEGs was similar in both species but still was 40% lower in the giant mole-rat than in the rat. Across tissues this summarizes to significantly less DEGs during aging in the giant mole-rat in comparison to the rat (p=0.016, Wilcoxon signed-rank test).

**Figure 1.**
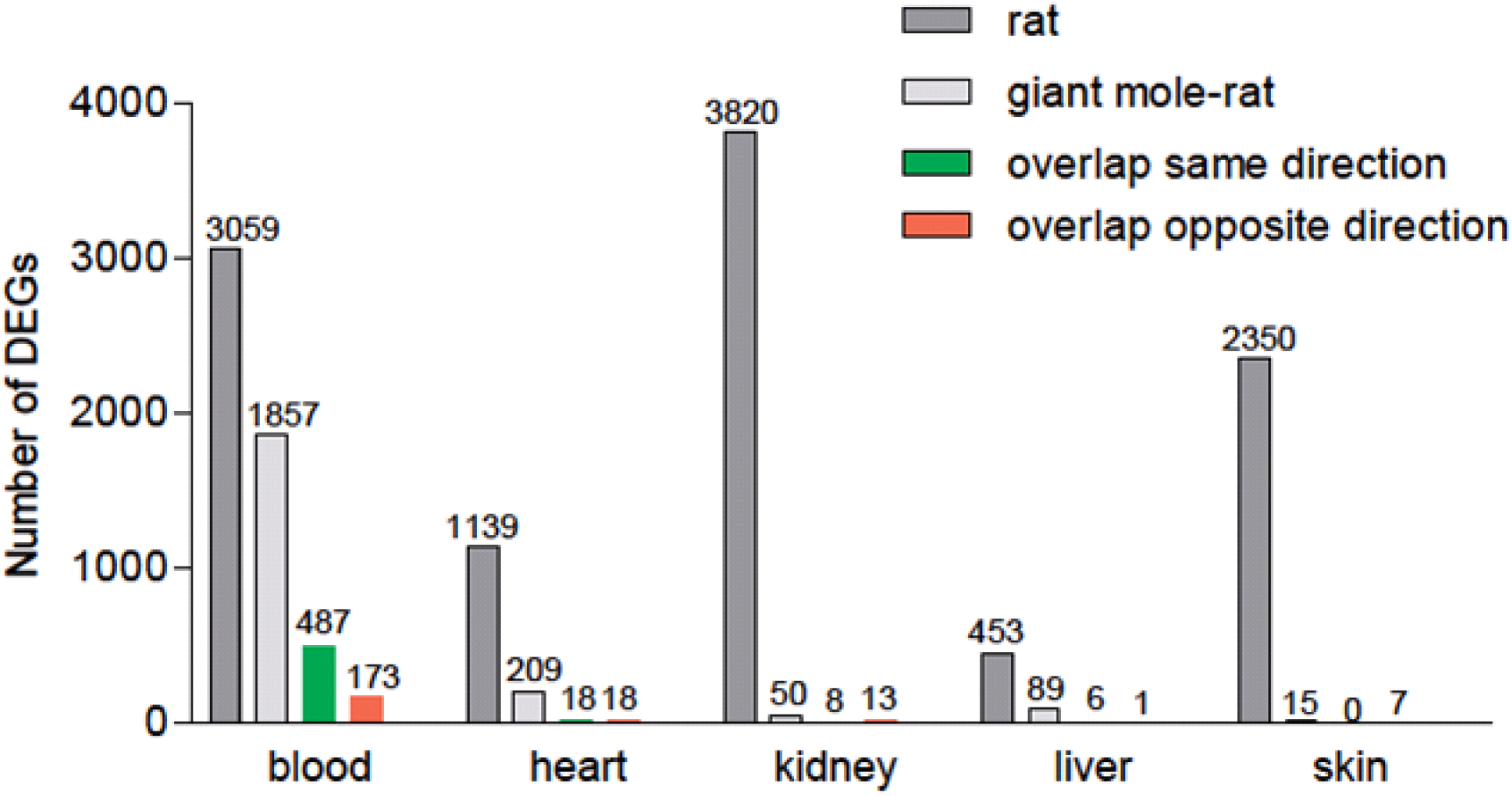
Numbers of differentially expressed genes (DEGs) during aging in five tissues of rat (*R. norvegicus*) and giant mole-rat (*F. mechowii*). Only orthologous genes in both transcript catalogs were counted.

This transcriptome stability of giant mole-rats during aging concurs with a general pattern of stability that has emerged from numerous molecular and physiological comparisons of the extremely long-lived naked mole-rat (*Heterocephalus glaber*, a close relative of giant mole-rats) with shorter-lived species mice or rats. For example, naked mole-rats maintain an unchanged membrane lipid composition during aging [3], a fairly stable production of reactive oxygen species [10] and relatively stable levels of oxidative damage on lipids [2], as well as high protein stability and integrity [11]. At the same time, all these parameters, which are known to be among the key factors for lifespan and age-related diseases [12], changed significantly in the unfavorable direction during aging in short-lived mice or rats. Naked mole-rats also show minimal decline of physiological functions, a maintenance of activity, fertility and body composition into old age, a remarkable resistance to cancer as well as mortality rates that do not increase obviously with age [1]. Given the close relatedness of naked and giant mole-rats and our own husbandry experience with the latter, we assume that several of the aforementioned properties are shared by both species.

Somewhat in line with our results, it has been reported earlier that gene expression in three naked mole-rat tissues remained nearly unchanged during the first half of lifespan [13]. However, this analysis had very limited statistical power as only one replicate per age was used. Regarding rats, our results are in good agreement with the rat body map initiative [14]. The database shows many DEGs – 491 to 12708 – across eleven tissues during rat aging using similar time points as we did (21 weeks vs. 2 years). The results of Kim *et al.* and the rat body map project cannot be directly compared since they used different methods for sequencing and DEG detection. Therefore, in this work we applied the same sequencing procedure as well the same bioinformatic analyses and confirmed that the transcriptomes of a long-lived African mole-rat species indeed remain stable during aging from young adulthood up to median lifespan in contrast to a short-lived rodent. Since gene expression is a basic regulatory process of the cell that underlies many of the above-mentioned molecular phenotypes and physiological observations, we suggest that transcriptome stability during aging is one of the key causal factors for the extraordinary long life‐ and healthspan of this, and maybe all, African mole-rat species.

Consistent with this idea, we found classical aging signatures across all examined tissues when looking at biological processes that were affected by differential gene expression in the rat, (Fig. 2). For instance, transcriptional alterations of “immune response” (GO:0006955, tables S12-21) and “inflammatory response” (GO: 0006954) genes are known as hallmarks of aging [15]. These processes, as well as many related processes such as response to cytokine (GO: 0034097) and leukocyte aggregation (GO: 0070486), are consistently enriched for DEGs in all examined rat tissues. In the giant mole-rat, on the other hand, we found these signatures only in blood. Summarizing the processes enriched for DEGs using REVIGO [16] results for all rat tissues and giant mole-rat blood in a largest summarized category that holds mainly those immune processes and is accordingly named “immune process” (blood, kidney and skin), “regulation of immune process” (heart) and “response to external stimulus” (liver) (Fig. S1-9). Other aging-relevant processes that are enriched for DEGs across rat tissues are, e.g., apoptotic process (GO: 0006915, all tissues except heart), coagulation (GO: 0050817, all tissues) and oxidation-reduction process (GO: 0055114, all tissues except liver). Again, these processes are enriched only in blood with regard to giant mole-rat DEGs. These results indicate that giant mole-rats evolved a slow-down of typical aging dependent transcriptional alterations in several vital tissues.

**Figure 2.**
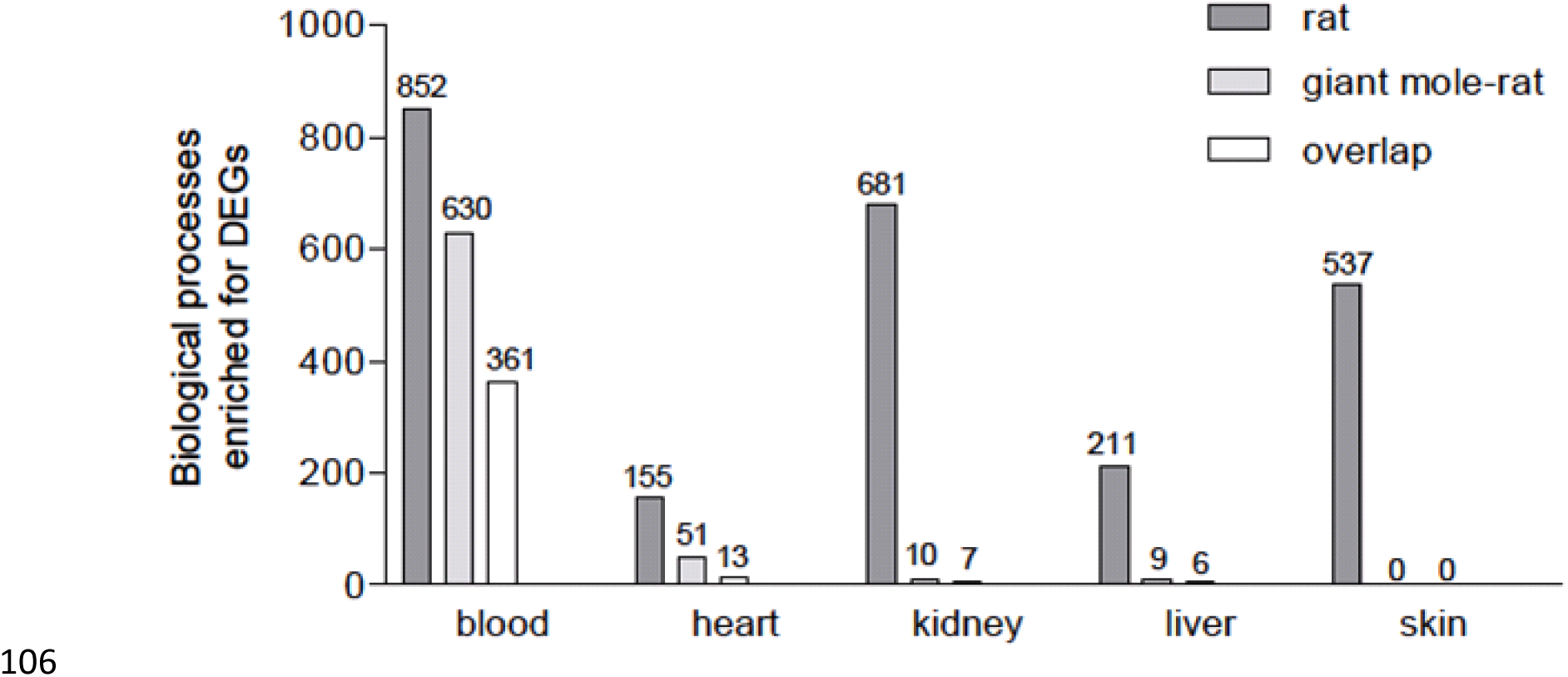
Numbers of biological processes (gene ontology) enriched for differentially expressed genes during aging in five tissues of rat (*R. norvegicus*) and giant mole-rat (*F. mechowii*).

On the single gene level, there is a modest but still significant (p <0.05, Fisher’s exact test) overlap between the DEGs of rat and giant mole-rat in blood, heart and skin as well as a tendency in kidney and liver (p <0.10) (Fig. 1, Tables S22-26). In the intersection of blood DEGs those are overrepresented that are regulated in the same direction during aging in both species (p=3.3*10^−31^, Fisher’s exact test based on regulation of all genes), fitting the shared aging-signatures in this tissue (see above). Interestingly, we found on the contrary an overrepresentation of DEGs that are regulated in opposite directions in skin (p=0.005). This points to the intriguing possibility that in some tissues expression changes that cause aging in the rat are counteracted by opposite changes during aging in the giant mole-rat. Also in kidney the majority of shared DEGs is regulated in opposite directions during aging (Fig. 1). As an example, “collagen metabolic process” (GO: 0032963) is one of the seven processes that are enriched in the kidney both in rat and giant mole-rat. While the enrichment in the rat is based on 20 collagen genes that are significantly up‐ and one down-regulated during aging, in giant mole-rat it results from four collagens and two genes coding for potent collagenases (*CTSK* and *CTSS*, [17]) all being down-regulated during aging. Of the latter six genes, five overlap with those that are significantly up-regulated in rat. Collagen regulation in the rat reflects the molecular aging process because lowering collagen levels attenuates kidney diseases in rats [18], while increased collagen levels in kidney were shown to induce cyst development in polycystic kidney disease in this species [19]. At the same time kidney diseases are a major cause of death in rats [20] and potentially also in (naked) mole-rats [21, 22]. The opposite collagen regulation pattern in giant mole-rat can be interpreted as an anti-aging program rather than a signature of the aging process.

In conclusion, we hypothesize that the higher transcriptome stability observed in long-lived giant mole-rats compared to short-lived rats evolved under different evolutionary constraints and contributes to the considerably distinct life history traits in short‐ and long-lived species: early onset and fast aging on one side and delayed/slowed down aging from young to elderly adulthood on the other.

## Methods

### Experimental design

The transcriptomes of young versus old animals from two species ‐ Wistar rats (*R. norvegicus*) and giant mole-rats (*F. mechowii*) ‐ were compared in this study. Five tissues (blood, heart, kidney, liver and skin) were sampled from both species and age cohorts. All examined animals were non-breeding males. Young rats had an age of 6 months, and old rats of 2 years. Young mole-rats had an age of 1.3-2 (mean 1.53) years, and old mole-rats of 5.5-7.7 years. The number of biological replicates per tissue for each age cohort and species was 4-8 depending on the tissue (Table S1/S27). All animals were housed and euthanized compliant with national and state regulations.

### Transcript catalogue sequences

The assembly of the giant mole-rat transcript catalog was performed based on recently published read data ([23], ENA study PRJEB20584) and the assembly framework FRAMA [24] using default parameters. For rat, mRNA sequences were obtained from RefSeq. For both species, only the longest transcript isoform per gene was used resulting in 15,864 and 23,479 reference transcripts/genes for giant mole-rat and rat, respectively.

### RNA-seq, read mapping and quantification

Tissue samples were collected and stored in RNA later (Qiagen), following isolation. Purification of RNA, for all tissues except blood, was done using Qiagen RNeasy Mini Kit following the manufacturer’s protocol. Blood samples (100 μl) were collected in RNAprotect Animal Blood reagent (Qiagen). The resulting RNA was purified RNeasy Protect Animal Blood Kit (Qiagen). Kidney and heart samples were treated with proteinase K before extraction, as recommended by the manufacturer. Poly(A) selection and preparation of the RNA-seq libraries was done using the TruSeq RNA v2 kit (Illumina). RNA-seq was performed using single-end sequencing with 51 base pairs on an Illumina HiSeq 2500 sequencing device and with at least 17 mio. reads per sample as described in Table S27. The reads were aligned to the respective reference using the “aln” algorithm of the Burrows-Wheeler Alignment tool (BWA) [25] allowing no gaps and a maximum of two mismatches in the alignment. Only those reads were used for quantification that could be uniquely mapped to the respective gene.

Read data for rat and giant mole-rat were deposited as ENA study PRJEB23955 (Table S27).

### Differential expression analysis

The differential expression analysis was performed using DeSeq2 [26]. In both species, the old animals were compared against the young ones. Genes with a p-value < 0.05 after correcting for multiple testing with the Benjamani-Hochberg method were considered as differentially expressed (Tables S2-S11). Biological processes that were enriched for DEGs were determined using gene ontology (GO, annotation package: org.Hs.eg.db) categories and Fisher’s exact test. Resulting pvalues were corrected for multiple testing with the Benjamini-Hochberg method. Additionally, GO categories with a p-value < 0.05 after correcting for multiple testing were summarized using REVIGO (cutoff=0.70, measure=SimRel, database=whole Uniprot) [16] (Fig. S1-9).

## Supporting Information listing

**Table S1.** Overview of examined animals.

**Table S2-S11.** Result of DESeq2-analysis for differentially expressed genes during aging in rat and giant mole-rat (one table per species and tissue).

**Table S12-S21.** Biological process gene ontologies that are enriched for DEGs (FDR <0.05) in rat and giant mole-rat (one table per species and tissue).

**Table S22-S26.** Overlap of genes that are differentially expressed in rat and naked mole-rat blood (one table per tissue).

**Table S27.** Samples that were sequenced in this study.

Figure S1-S9. REVIGO treemap of gene ontology processes that are significantly enriched (FDR <0.05) for gene ontology processes (one figure for tissue and species, giant mole-rat skin is missing because the number of enriched terms was too small for summarization).

## Acknowledgements

We thank Ivonne Görlich, Petra Dobermann, Sabine Bischoff and Christoph Bergmeier for excellent assistance in the preparation of biological samples.

## References

1. Edrey, Y.H., et al., Successful aging and sustained good health in the naked mole rat: a long-lived mammalian model for biogerontology and biomedical research. ILAR J, 2011. 52(1): p. 41–53.

2. Andziak, B. and R. Buffenstein, Disparate patterns of age-related changes in lipid peroxidation in long-lived naked mole-rats and shorter-lived mice. Aging Cell, 2006. 5(6): p. 525–32.

3. Hulbert, A.J., S.C. Faulks, and R. Buffenstein, Oxidation-resistant membrane phospholipids can explain longevity differences among the longest-living rodents and similarly-sized mice. J Gerontol A Biol Sci Med Sci, 2006. 61(10): p. 1009–18.

4. Dammann, P., Slow aging in mammals-Lessons from African mole-rats and bats. Semin Cell Dev Biol, 2017. 70: p. 154–163.

5. Fushan, A.A., et al., Gene expression defines natural changes in mammalian lifespan. Aging Cell, 2015. 14(3): p. 352–65.

6. Malik, A., et al., Genome maintenance and bioenergetics of the long-lived hypoxia-tolerant and cancer-resistant blind mole rat, Spalax: a cross-species analysis of brain transcriptome. Sci Rep, 2016. 6: p. 38624.

7. Tacutu, R., et al., Human Ageing Genomic Resources: integrated databases and tools for the biology and genetics of ageing. Nucleic Acids Res, 2013. 41(Database issue): p. D1027–33.

8. Dammann, P., et al., Extended longevity of reproductives appears to be common in Fukomys mole-rats (Rodentia, Bathyergidae). PLoS One, 2011. 6(4): p. e18757.

9. Carlus, M., et al., Historical control data of neoplastic lesions in the Wistar Hannover Rat among eight 2-year carcinogenicity studies. Exp Toxicol Pathol, 2013. 65(3): p. 243–53.

10. Csiszar, A., et al., Vascular aging in the longest-living rodent, the naked mole rat. Am J Physiol Heart Circ Physiol, 2007. 293(2): p. H919–27.

11. Perez, V.I., et al., Protein stability and resistance to oxidative stress are determinants of longevity in the longest-living rodent, the naked mole-rat. Proc Natl Acad Sci U S A, 2009. 106(9): p. 3059–64.

12. Lopez-Otin, C., et al., The hallmarks of aging. Cell, 2013. 153(6): p. 1194–217.

13. Kim, E.B., et al., Genome sequencing reveals insights into physiology and longevity of the naked mole rat. Nature, 2011. 479(7372): p. 223–7.

14. Yu, C., et al., RNA sequencing reveals differential expression of mitochondrial and oxidation reduction genes in the long-lived naked mole-rat when compared to mice. PLoS One, 2011. 6(11): p. e26729.

15. de Magalhaes, J.P., J. Curado, and G.M. Church, Meta-analysis of age-related gene expression profiles identifies common signatures of aging. Bioinformatics, 2009. 25(7): p. 875–81.

16. Supek, F., et al., REVIGO summarizes and visualizes long lists of gene ontology terms. PLoS One, 2011. 6(7): p. e21800.

17. Barry, Z.T. and M.O. Platt, Cathepsin S cannibalism of cathepsin K as a mechanism to reduce type I collagen degradation. J Biol Chem, 2012. 287(33): p. 27723–30.

18. Liu, B., et al., Increasing extracellular matrix collagen level and MMP activity induces cyst development in polycystic kidney disease. BMC Nephrol, 2012. 13: p. 109.

19. Gilbert, R.E., et al., A purpose-synthesised anti-fibrotic agent attenuates experimental kidney diseases in the rat. PLoS One, 2012. 7(10): p. e47160.

20. Ettlin, R.A., P. Stirnimann, and D.E. Prentice, Causes of death in rodent toxicity and carcinogenicity studies. Toxicol Pathol, 1994. 22(2): p. 165–78.

21. Delaney, M.A., et al., Spontaneous histologic lesions of the adult naked mole rat (Heterocephalus glaber): a retrospective survey of lesions in a zoo population. Vet Pathol, 2013. 50(4): p. 607–21.

22. Delaney, M.A., M.J. Kinsel, and P.M. Treuting, Renal Pathology in a Nontraditional Aging Model: The Naked Mole-Rat (Heterocephalus glaber). Vet Pathol, 2016. 53(2): p. 493–503.

23. Sahm, A., et al., Long-lived rodents reveal signatures of positive selection in genes associated with lifespan and eusociality. bioRxiv, 2017.

24. Bens, M., et al., FRAMA: from RNA-seq data to annotated mRNA assemblies. BMC Genomics, 2016. 17: p. 54.

25. Li, H. and R. Durbin, Fast and accurate short read alignment with Burrows-Wheeler transform. Bioinformatics, 2009. 25(14): p. 1754–60.

26. Love, M.I., W. Huber, and S. Anders, Moderated estimation of fold change and dispersion for RNA-seq data with DESeq2. Genome Biol, 2014. 15(12): p. 550.

